# Investigation into the effects of sodium valproate on prepubertal mouse gonads *in vivo*

**DOI:** 10.64898/2026.09.01.748280

**Authors:** Adomas Liugaila, Rod T Mitchell, Adam Gadd, Kathleen Duffin, Agnes Stefansdottir

## Abstract

**Objective:** Sodium valproate (SV) is a widely used anti-epileptic drug with well-established reproductive and teratogenic effects, yet its impact on the developing prepubertal reproductive system remains poorly understood. This study investigated the effects of prepubertal SV exposure on gonadal development, including folliculogenesis, testicular architecture and steroidogenic gene expression in male and female mice *in vivo*.

**Methods:** CD1 mouse pups received intraperitoneal injections of saline (control), low or high dose SV (50 or 100 mg/kg, respectively) on postnatal days (PND) 6, 8, and 10. On PND17 the animals were culled and gonads dissected. Ovaries and testes were assessed histologically using haematoxylin and eosin staining. Ovarian follicle number, stage and health were quantified. Testicular tubule morphology and key testicular cell numbers (germ, Sertoli, spermatogonial stem cells, and interstitial/Leydig cells) were evaluated using immunofluorescence with automated image analysis. Expression of key steroidogenic genes (CYP11A1, STAR, CYP19A1 in ovaries; INSL3, STAR, CYP11A1 in testes) was measured by RT-qPCR.

**Results:** SV exposure did not significantly affect ovarian follicle number, distribution or health. Testicular morphology and the density of Sertoli, spermatogonial stem cells, and interstitial cells were likewise unaffected. However, a dose-dependent trend toward reduced germ cell density was observed in males at a high dose SV, though this did not reach statistical significance (P = 0.058). Steroidogenic gene expression was unaffected by SV exposure in both ovaries and testes across all treatment groups.

**Significance:** This study provides the first *in vivo* assessment of short-term prepubertal SV exposure on gonadal development in both sexes. The overall findings suggest that brief SV exposure during the prepubertal window does not cause overt gonadotoxicity. Nonetheless, the trend toward reduced testicular germ cell density warrants further investigation with longer exposure durations and functional fertility endpoints, to better inform clinical risk assessment in paediatric patients receiving SV.

## INTRODUCTION

Epilepsy is a neurological disorder characterised by recurrent unprovoked seizures caused by abnormal electrical activity in the brain and is the most common neurological condition in children, affecting between 41–187 per 100,000 children (1). If untreated or poorly controlled, seizures can result in permanent brain injury, mental health complications and sudden death (2). Management typically requires long-term treatment with anti-epileptic drugs (AEDs) such as sodium valproate (SV), which has been widely prescribed since the 1970s due to its strong efficacy and favourable pharmacoeconomic profile (3).

A growing body of evidence indicates that SV exerts adverse side effects on adults and children, as well as teratogenic effects in the exposed fetus (3–8). SV has been linked to endocrine disturbances affecting the hypothalamic-pituitary-gonadal (HPG) axis and reproductive function (9–13). In adult women, SV exposure has been associated with polyendocrine metabolic ovarian syndrome (PMOS), previously known as polycystic ovary syndrome (PCOS), which includes-hyperandrogenism and menstrual irregularities. In males, it has been linked to reductions in sperm count and motility, altered sperm morphology, and decreased gonadotropin levels (11, 14–18). SV has also been shown to disrupt hormone production and germ cell development in the human fetal testis, demonstrating endocrine-disrupting effects during reproductive development (19).

Growing concerns regarding the risks of SV have prompted progressive regulatory tightening. Following the introduction of the Pregnancy Prevention Programme (PPP), SV was contraindicated in pregnant women and women of reproductive age. In 2024, the Medicines Healthcare Regulatory Agency (MHRA) mandated that SV prescriptions for all patients under 55 years require authorisation from two independent specialists, confirming the absence of suitable alternative treatment options (20). Prior to implementation of these restrictions, considerable numbers of patients, including children, received SV, and it remains the most effective treatment option for a subset of individuals. Characterising the long-term reproductive consequences of SV exposure is therefore of considerable clinical importance.

The reproductive systems develop from fetal life through to puberty. In the ovary, the primordial follicle pool is established during fetal development and consists of oocytes surrounded by granulosa cells, with theca cells differentiating as follicles grow and with granulosa-theca cell interactions supporting steroidogenesis (Figure 1A). During the prepubertal period, follicles typically develop only to early growing or pre-antral stages due to limited gonadotropin stimulation (21). Following activation of the HPG axis in puberty, follicular development starts progressing to antral and pre-ovulatory stages, culminating in ovulation (22; Figure 1A). The testes contain Leydig cells, which produce testosterone, Sertoli cells, which support germ cell development, and germ cells, which give rise to spermatogonia and eventually spermatozoa from puberty onwards (Figure 1B). During the prepubertal period, although spermatogenesis is not yet established, important developmental changes occur. Sertoli cells undergo maturation, germ cells proliferate at a low rate, and although the fetal Leydig cells are largely quiescent, they go through a transient neonatal period of testosterone production, known as ‘mini-puberty’, before being replaced by the adult Leydig cell population (23).

**Figure 1.**
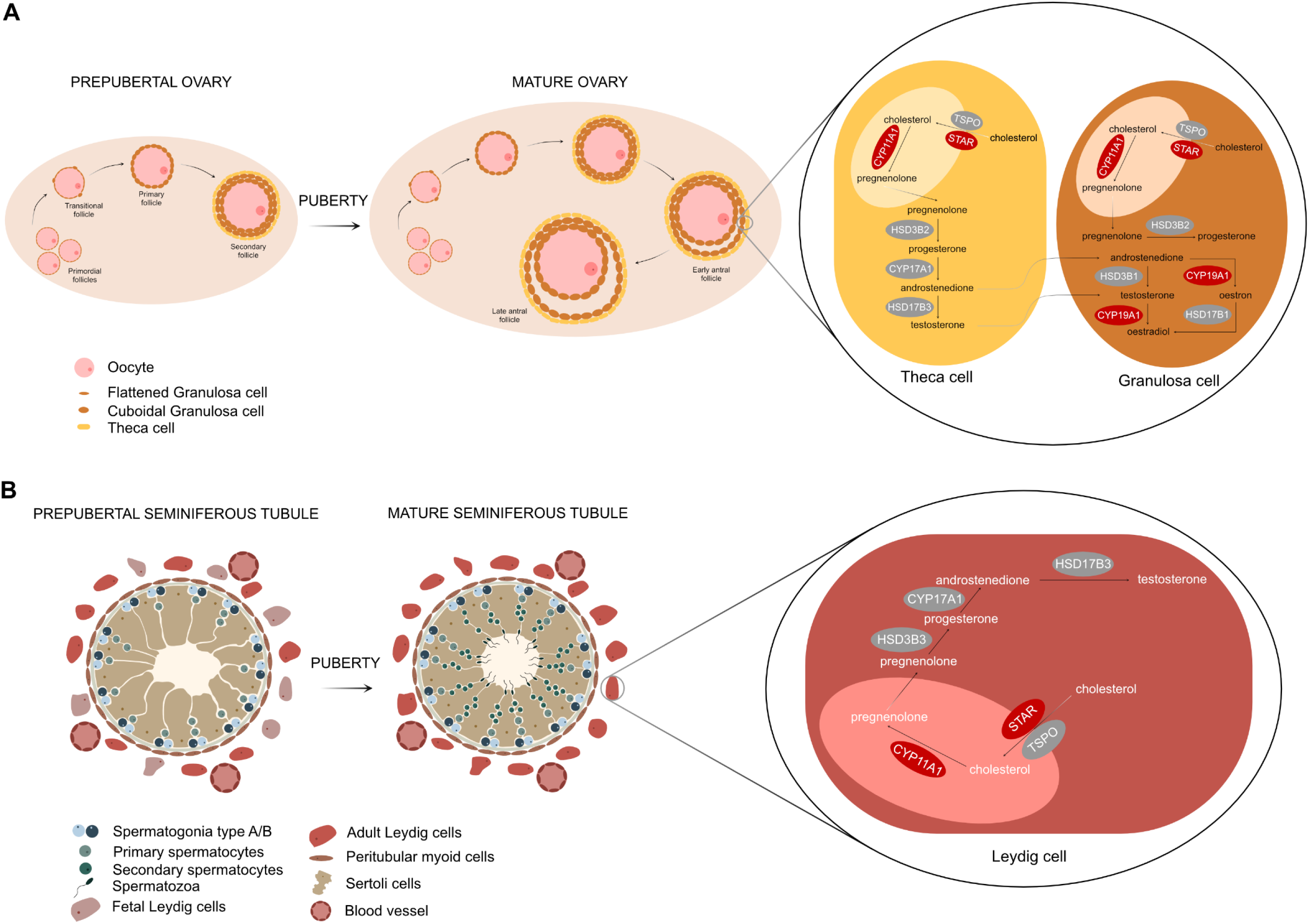
(A) Follicle development in the prepubertal and mature ovary & simplified oestradiol and progesterone production pathway in theca and granulosa cells in the ovary. Mature ovary contains primordial, transitional, primary, secondary and antral follicles. Mature oocytes are only present after puberty, when HPG axis is activated. (B) Differences in the development of prepubertal and mature seminiferous tubules & testosterone production pathway in Leydig cells. A prepubertal seminiferous tubule is made up of peritubular myoid (PTM), Sertoli- and Leydig cells, as well as spermatogonia and sporadically appearing primary spermatocytes. While a mature seminiferous tubule also contains secondary spermatocytes and mature spermatozoa. Genes encoding proteins in the steroidogenic pathway were investigated in our study using RT-qPCR are noted in maroon. Figure created using Affinity.

The prepubertal gonadal developmental window is sensitive to endocrine disruption. Exposures to drugs, including SV, may impair gonadal development, folliculogenesis, and/or steroid hormone production, with potentially lasting consequences for fertility. Previous research has focused predominantly on SV’s reproductive effects in adults, with considerably less known regarding its impact on gonadal development in children, whose reproductive system differs substantially from that of adults. These include differences in structure as well as cellular and hormonal activity; immature gonads may exhibit heightened or otherwise distinct sensitivity to environmental exposures during critical developmental windows. It is therefore important to evaluate the effects of SV on prepubertal reproductive health.

## MATERIALS & METHODS

### Animals and experimental procedure

All experiments were approved by the University of Edinburgh’s Local Ethical Review Committee and carried out in accordance with UK Home Office regulations under the Animals (Scientific Procedures) act (ASPA) 1986. Wild-type CD-1 mice were maintained and bred in an environmentally-controlled room on a 14-h light:10-h dark photoperiod.

A feasibility study was performed trialling doses of 50, 100 and 500 mg/kg SV and based on tolerable end points, where 100 mg/kg was determined as the safe upper limit. Mouse pups received three intraperitoneal injections on postnatal days (PND) 6, 8 and 10 of either saline, a low dose of 50 mg/kg (LD), or a high dose of 100 mg/kg (HD) of valproic acid sodium salt (Merck Life Science; P4543-10g) diluted in saline solution. The pups were culled on PND17. Female and male pups were weighed at the time of culling, and testis weight measured.

### Freezing, fixation and processing of gonads

Ovaries and testes were placed in 1X phosphate-buffered saline (PBS; Fisher Scientific). The right gonads from both sexes were placed into an Eppendorf tube and snap-frozen in a dry-ice ethanol bath. The left ovaries were fixed in 10% neutral-buffered formalin solution (NBF, Sigma Aldrich Ltd) for 24 hours at room temperature. The left testis was halved: one half was fixed in Bouin’s fixative (Sigma Aldrich Ltd) and the other half in 10% NBF solution (Sigma Aldrich Ltd) for 24 hours at room temperature. After fixation, all tissue was washed in 70% ethanol, processed and embedded in wax. Wax blocks were sectioned using a microtome (Leica RM2255) at 5μm thickness and placed on microscope slides.

### Histological assessment of ovaries

To assess follicular number and health in the ovaries, every tenth section of ovary was stained using haematoxylin & eosin (H&E) staining and then imaged using an inverted brightfield microscope at 20X (ECLIPSE Ti2; Nikon Instruments). Follicle counts and assessment of follicle stage and health were performed in a blinded manner using Fiji software (24). Follicles were counted only if an oocyte contained a germinal vesicle. Follicles comprised of a flattened granulosa cell layer were considered primordial follicles (PMFs), those containing flattened and cuboidal granulosa cells were assigned as transitional follicles (TRNS). Follicles containing a single or double layer of cuboidal granulosa cells were classed as primary (PRIM) or secondary (SEC) follicles, respectively. If a follicle had a small antrum starting to form, it was assigned as early antral (E ANTRAL), or if it had a fully formed large antrum, it was considered as late antral (L ANTRAL). Follicles consisting of a round, centrally placed oocyte with an evenly stained nucleus and healthy non-pyknotic granulosa cells were assigned as healthy, while oocytes with shrunken and pyknotic nucleus and/or granulosa cells were considered unhealthy (25).

### Histological assessment of testes

To histologically assess testicular health, three H&E-stained sections separated by twenty intervening sections (≥ 100µm apart) were imaged for each testis using an inverted brightfield microscope at 20X (ECLIPSE Ti2; Nikon Instruments). H&E-stained testes sections were analysed using Fiji software with the assessor blind to treatment. Only round tubules with a clear lumen were analysed. Round tubules were identified by measuring two perpendicular diameters of each tubule; if the ratio between these diameters was between 0.8 and 1.2, the tubule was classified as round. This was set in order to ensure that only the tubules that had been cut transversely were included. Moreover, only tubules with a clear lumen were analysed. Area of the whole tubule, its germinal epithelium and lumen was acquired. Furthermore, the depth of the germinal epithelium was quantified by calculating the mean of the narrowest and the widest points of the tubules were measured and their mean calculated. Ratios of germinal epithelium area to tubular area, lumen area to tubular area and lumen area to germinal epithelium area were then quantified.

### Immunofluorescence

Immunofluorescence was carried out to identify the density of germ-, interstitial-, Sertoli- and spermatogonial stem (SSCs) cells in the testis. Vasa homologue (MVH) was used as a germ cell marker, COUP transcription factor II (COUPTFII) as a marker for interstitial cells (including Leydig cells), SRY-Box Transcription Factor 9 (SOX9) as a marker for Sertoli cells and Promyelocytic Leukaemia Zinc Finger protein (PLZF) as a marker for SSCs (Table S1).

Dewaxing of slides was carried out in xylene followed by rehydration of the sections. Antigen retrieval was performed in 10mM citrate buffer (pH 6, Sigma Aldrich), followed by a wash in 1X PBS. Tissue sections were covered in a blocking solution (20% goat serum, 5% bovine serum albumin (Sigma Aldrich), 1X PBST (1X PBS, 0.1% Triton-X)) for 1 hour at room temperature in a humidified environment. Slides were incubated with primary antibodies diluted in the blocking solution overnight at 4°C in a humidified environment, which was followed by incubation with secondary antibody for one hour the next day (Table S1). Two negative control slides were included for each of the immunofluorescence experiments: one without primary antibodies and one without secondary antibodies. All slides were submerged in 4’,6-diamidino-2-phenylindole (DAPI; Invitrogen) diluted 1:5,000 in 1X PBS for 5 minutes to counterstain the nuclei. Vectashield (Vector Laboratories) was applied and the slides were coverslipped. Three sections separated by ten intervening sections were imaged using an inverted microscope at 20X magnification (ECLIPSE Ti2; Nikon Instruments).

### Image analysis

Images of immunofluorescence were analysed using QuPath software (26) in an automated and blind manner (27). Firstly, regions of interest (ROIs) were segmented using a multilayer perceptron artificial neural network (ANN-MLP). Models were trained on all available fluorescent channels and on images spanning experimental groups, antibody controls and vehicle using selected features; Gaussian filter (intensity and colour), gradient magnitude (edges), structure tensor coherence (cell orientation) and Hessian determinant (blob-like structures). Within each ROI, individual cells were segmented using a CNN-based star-convex polygon approach (StarDist; https://github.com/s/tardist/stardist; DOI:10.48550/arXiv.1806.03535). This enabled accurate identification of nuclei and reconstruction of full cell boundaries, including cytoplasmic regions. The cells were then classified by fluorescence location as well as intensity patterns. Cells were categorised according to marker-specific thresholds, e.g. nuclear intensity of PLZF or cytoplasmic intensity of MVH. Cells not meeting classification criteria were designated as unclassified. Settings determining the threshold of the signal were kept the same for each immunohistochemistry run. Following classification, cell counts, total area of tubules or interstitial tissue area were calculated. Germ-, Sertoli- and spermatogonial stem cell densities per tubule area were acquired, as well as interstitial cells density per interstitial area.

### Quantitative real-time polymerase chain reactions

In order to examine gene expression of key steroidogenic enzymes in the ovaries and testes, RNA was extracted from tissue lysate using the RNeasy Micro Kit (Qiagen), according to manufacturer’s instructions. To remove genomic DNA contamination, RQ1 RNase-Free DNase treatment was performed (Promega), according to manufacturer’s instructions. The concentration of the RNA was measured using NanoDrop One (ThermoFisher). A 260/280 optical density ≥2 indicated that the RNA was suitably pure for further use. Five hundred ng of template RNA were used to synthesise cDNA using Maxima first strand cDNA synthesis kit (ThermoFisher), as per manufacturer’s instructions. Negative controls (RT-) were prepared using identical reactions where the Maxima enzyme mix was replaced with nuclease-free dH_2_O.

Primer specifications used in the study can be seen in Supplementary Table 2. Efficient primers were used for further RT-qPCRs to establish fold change in gene expression. For further RT-qPCRs, cDNA of each sample was run in 1:4 dilution using the same KAPA SYBR FAST qPCR kit (Roche), where RT- was included for every sample. qPCRs were run on the same protocol as for primer efficiency, according to manufacturer’s instructions. Fold-change in mRNA expression levels was calculated.

### Statistical analysis

Statistical analyses were performed using GraphPad Prism (GraphPad Software). Kolmogorov-Smirnoff tests were conducted to assess data normality. If data was distributed normally, one-way ANOVA was performed, followed by the Bonferroni post-hoc test to determine statistical significance between control and treatment groups. Kruskal-Wallis non-parametric test was used, where data was not normally distributed. Results were considered statistically significant where p < 0.05.

## RESULTS

### Prepubertal exposure to SV does not affect female or male pup weight

The impact of SV treatment on pup and gonadal weight was examined. No significant differences were found in body weight between PND6 and PND17 in either female (Figure S1, P = 0.16) or male (Figure S1, P = 0.68) pups. In addition, the testis weight to body weight ratio remained unchanged (Figure S1, P = 0.11).

### Prepubertal SV exposure does not affect ovarian follicle number, distribution or health

To examine the effect of SV treatment on the prepubertal ovaries, ovaries were histologically analysed to assess for follicle number, distribution and health (Figure 2A). There was no significant difference in the total number of follicles between the control group and those exposed to either the low or high dose of SV (Figure 2B, P = 0.89). When follicles were further categorised into different types, there were no significant differences in follicle distribution between controls and the two treatment groups (Figure 2C, P > 0.05). Furthermore, no differences were observed in ratio of healthy to unhealthy follicles, total number of unhealthy follicles, or distribution of follicle types (Figure S2).

**Figure 2.**
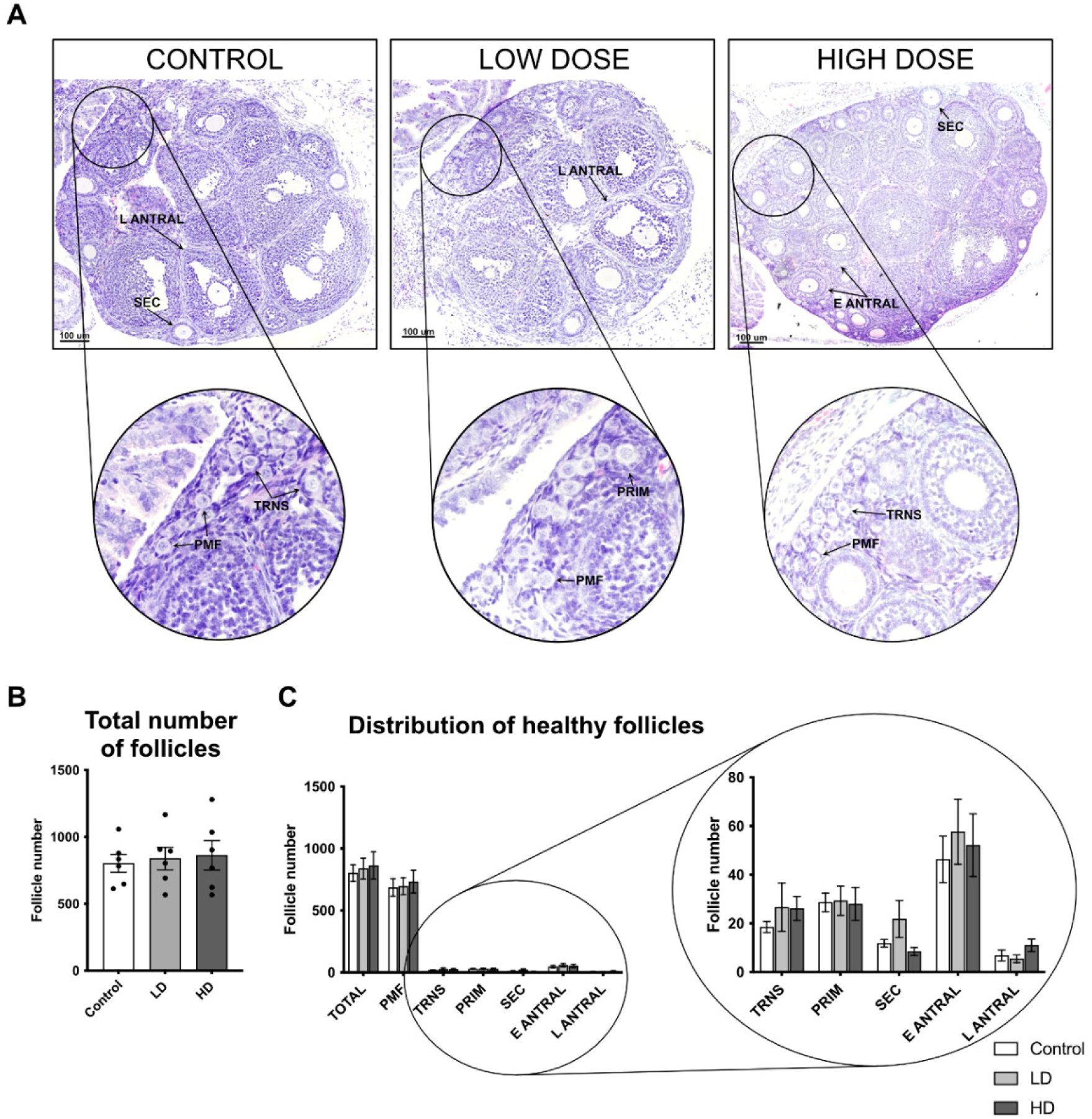
Representative images of H&E-stained ovaries treated with control (saline), low dose (LD: 50 mg/kg) or high dose (HD: 100 mg/kg) of SV (A). Primordial (PMF), transitional (TRNS), primary (PRIM), secondary (SEC), early antral (E ANTRAL) and late antral (L ANTRAL) follicles. Exposure to SV does not affect the total number of healthy follicles (B), or their distribution (C) in the prepubertal mouse ovaries at PND17. Bars represent mean ± SEM; n = 6 for each group. Figure created using Affinity.

### Prepubertal SV exposure does not affect testicular morphology

Next, histological examination of the testes was undertaken to examine whether SV treatment could disrupt testicular architecture (Figure 3A). A quantitative H&E analysis was performed on the testicular tissue sections, where the ratio of germinal epithelium area to tubular area and the ratio of lumen area to tubular area found to be unchanged between groups (P = 0.44, Figure 3B). Moreover, there were no significant differences in the depth of germinal epithelium across the different treatment groups (Figure 3C, P = 0.42), and the ratio of tubular to interstitial area remained unchanged (Figure 3D).

**Figure 3.**
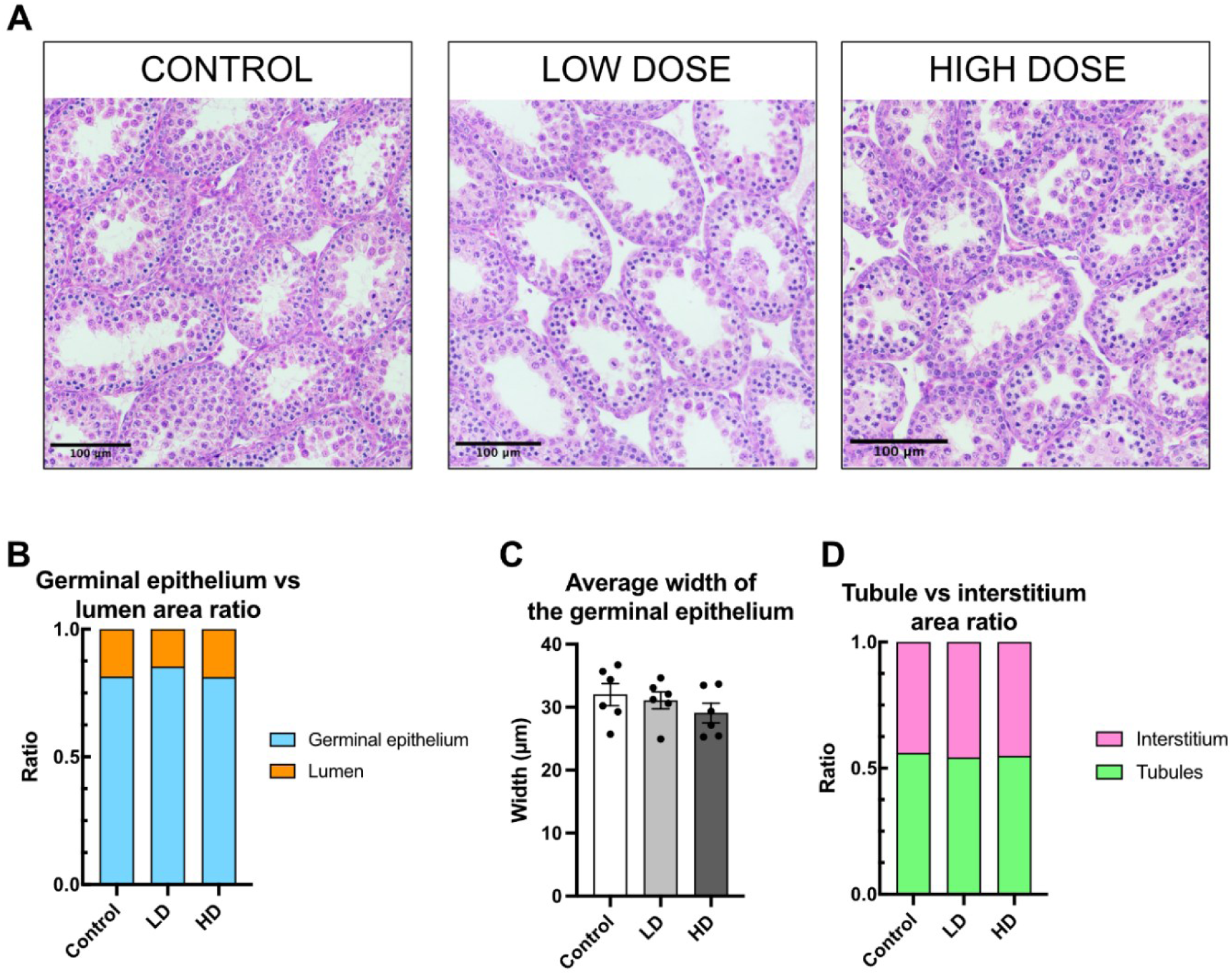
Representative H&E images of prepubertal mouse testes at PND17 exposed to control (saline), low dose (LD: 50 mg/kg) or high dose (HD: 100 mg/kg) of SV (A). Scale bar indicates 100 μm. Effect of SV on germinal epithelium vs lumen area (B), germinal epithelium depth (C) and interstitial vs tubular area (D). No significant differences were observed between groups. Bars represent mean ± SEM; n = 6 per group. Figure created using Affinity.

### Prepubertal SV exposure does not affect key testicular cell types

To gain insight into any potential impact on key testicular cell types, immunohistochemical analysis was carried out on testicular tissue. SV exposure did not affect the number of germ or spermatogonial stem cells in the testes. Although a dose dependent trend towards a lower germ cell (MVH+) density in the HD group was observed (control: 4785.34 ± 413.8, vs HD 3202.39 ± 515.1; mean ± SEM), this did not reach statistical significance (Figures 4A,B, P = 0.058). The density of SSCs (MVH^+^:PLZF^+^) was unchanged between control, LD and HD (Figures 4A and C, P = 0.14).

**Figure 4.**
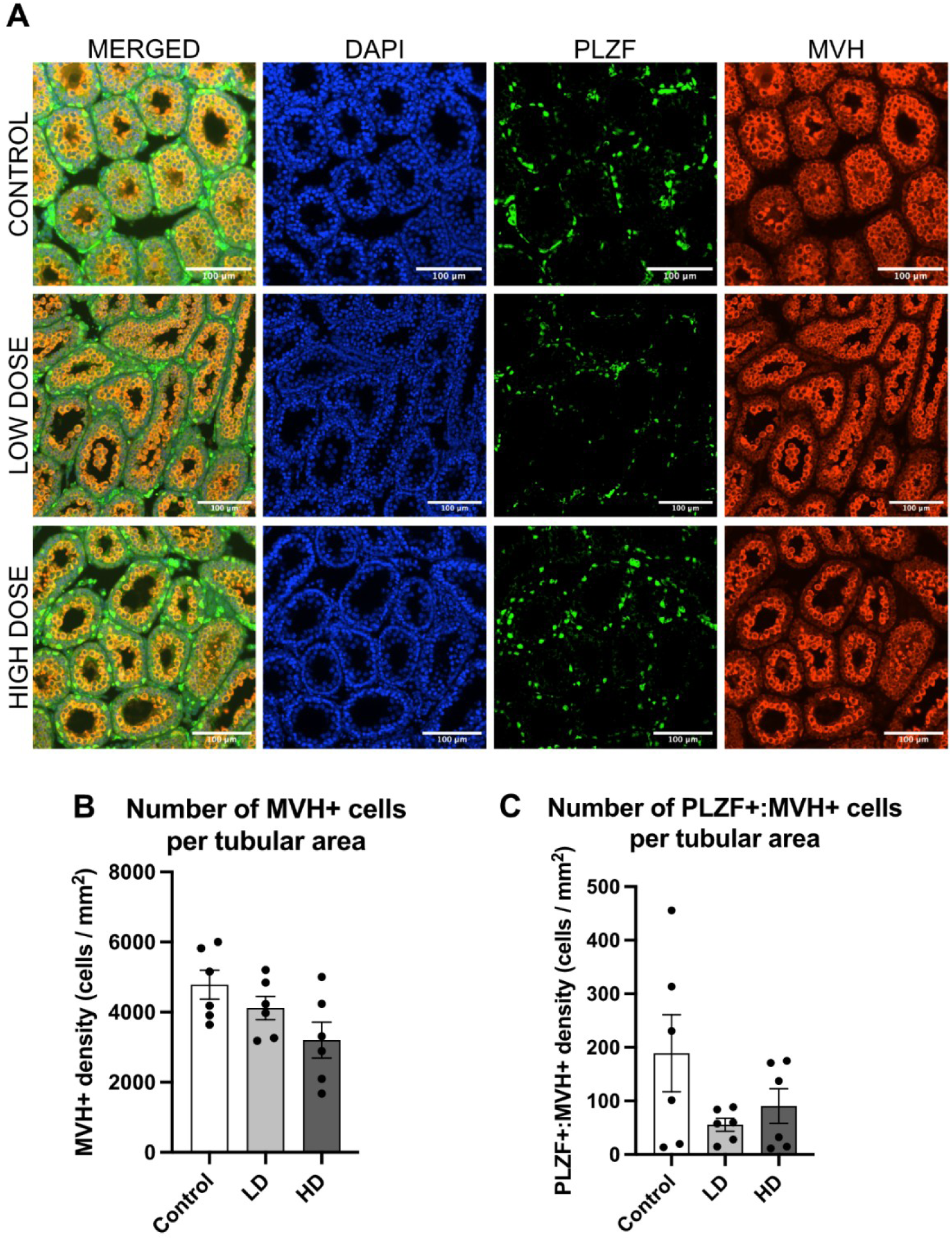
Representative images of prepubertal mouse testes at PND17 following exposure to control (saline), low dose (LD: 50 mg/kg) or high dose (HD: 100 mg/kg) of SV (A). Immunofluorescence staining for the spermatogonial stem cell marker PLZF (green) and the germ cell marker MVH (red), with DAPI nuclear counterstain (blue). Scale bar = 100 μm. MVH⁺ cell density per tubular area (B) and PLZF⁺:MVH⁺ cell density per tubular area (C). A dose-dependent trend toward reduced germ cell density was observed (P = 0.0538), while SSC density per tubule was unchanged. Bars represent mean ± SEM; n = 6 per group. Figure created using Affinity.

Sertoli cell density remained unchanged between control, LD and HD groups (Figures 5A and B, P = 0.18). There were also no significant differences in the density of interstitial, which includes Leydig cells (COUPTFII^+^) cells across different treatment groups (Figures 5C and D, P = 0.55).

**Figure 5.**
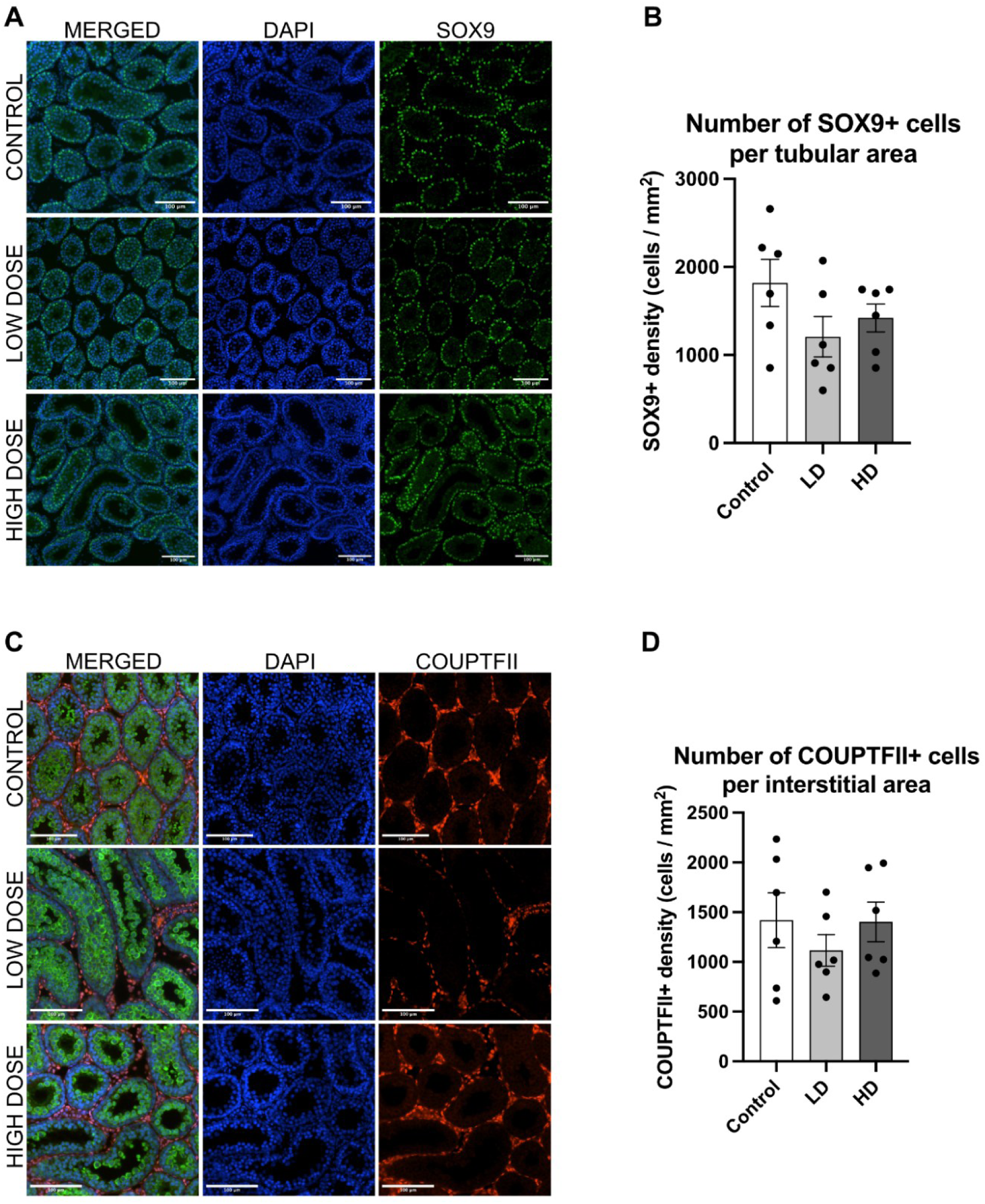
Representative images of prepubertal mouse testes at PND17 following exposure to control (saline), low dose (LD: 50 mg/kg) or high dose (HD: 100 mg/kg) of SV. Immunofluorescence staining for the Sertoli cell marker SOX9 (green) with DAPI nuclear counterstain (blue) (A), and Sertoli cell density per tubular area (B). Immunofluorescence staining for the germ cell marker MVH (green) and the interstitial cell marker COUPTFII (red) with DAPI (blue) (C), and COUPTFII⁺ cell density per interstitial area (D). Scale bar = 100 μm. No significant differences were observed between treatment groups. Bars represent mean ± SEM; n = 6 per group. Figure created using Affinity.

### Exposure to SV does not affect the expression of key steroidogenic genes in the prepubertal gonads

To evaluate the effect of SV exposure on the expression of key steroidogenic enzymes in the prepubertal gonads, qPCR was carried out to measure the expression of CYP11A1, STAR and CYP19A1 in the ovary (Figure 6A), and the expression of INSL3, STAR and CYP11A1 in the testis (Figure 6B). No significant differences were found in the expression of CYP11A1 (P = 0.09), STAR (P = 0.59) or CYP19A1 (P = 0.71) in the ovary, nor in INSL3 (P = 0.91), STAR (P = 0.59) or CYP11A1 (P = 0.75) in the testis between the treatment groups and controls.

**Figure 6.**
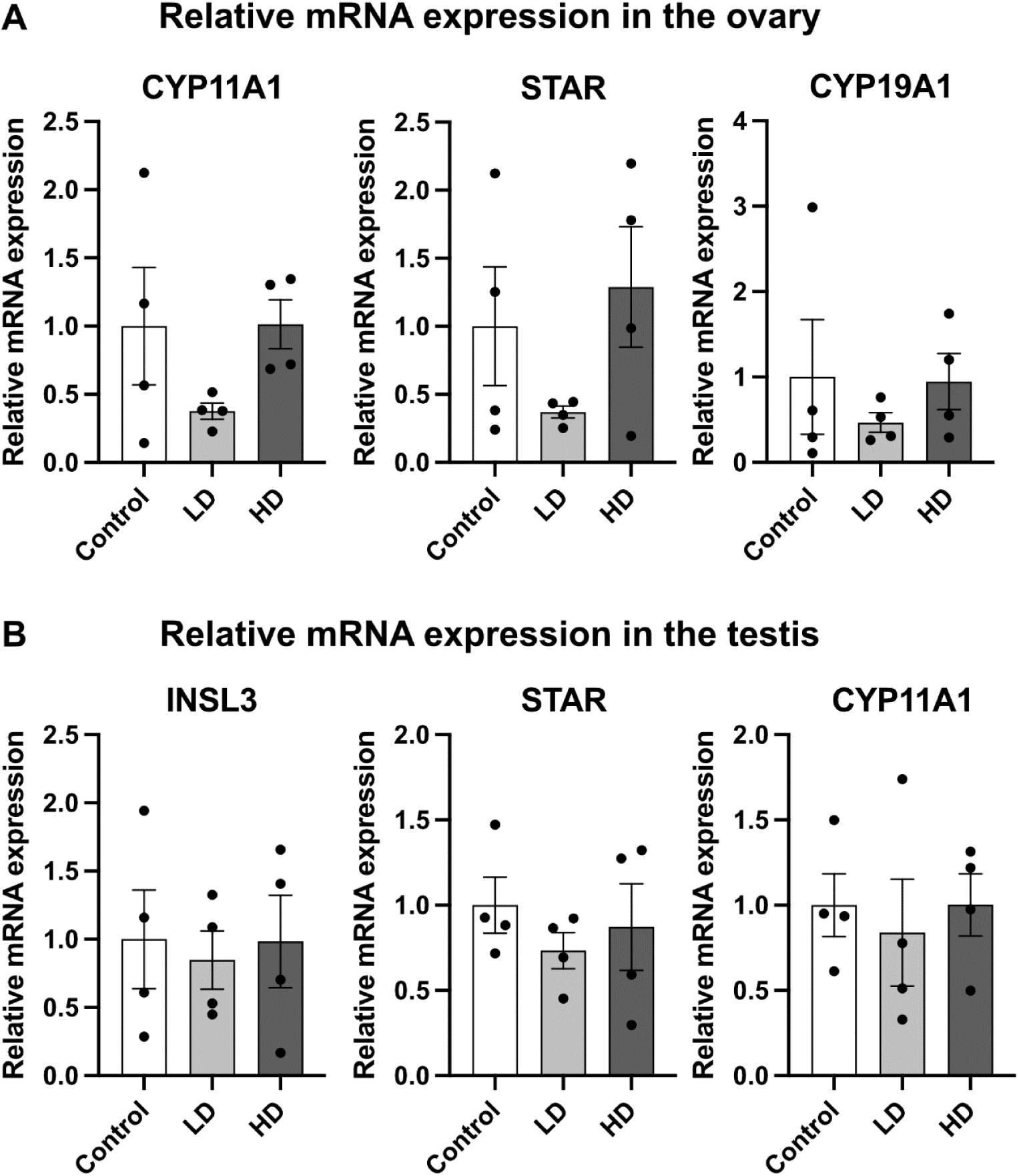
Exposure to SV does not affect the expression of steroidogenic genes in the ovary or testis. RT-qPCR analysis showed no significant differences between control, low dose (low dose (LD: 50 mg/kg) or high dose (HD: 100 mg/kg) groups in ovarian expression of CYP11A1, STAR, and CYP19A1 (A) or testicular expression of INSL3, STAR, and CYP11A1 (B). Bars represent mean ± SEM; n = 4 per group. Figure created using Affinity.

## DISCUSSION

SV is a widely prescribed and effective AED whose teratogenic effects have prompted increasingly stringent prescribing regulations. While the teratogenic risks of *in utero* exposure are well established, much less is known about the impact of SV on the developing reproductive system. Here, we used an *in vivo* mouse model to investigate the effect of SV treatment on prepubertal gonads. Neither dose of SV caused significant gonadotoxic effects on ovarian follicle number, distribution or health; testicular cell density; or steroidogenic gene expression in either gonad. A dose-dependent trend towards reduced testicular germ cell number was observed, but this did not reach statistical significance. These findings suggest that prepubertal SV exposure does not substantially compromise gonadal development in mice.

Females are born with a fixed number of oocytes that cannot be replenished (28). At the stage of development assessed herein, the follicle pool had already been established; however, the prepubertal period is an important stage for ovary maturation and endocrine regulation (29). Granulosa cells of growing follicles secrete AMH, which helps preserve the ovarian reserve by preventing premature activation of the dormant follicles (30). They also produce oestradiol, which controls development and selection of the dominant follicles, while also ensuring follicle survival and growth (31). Moreover, oestradiol triggers the activation of HPG axis during puberty, thereby establishing regular reproductive cycles (32). Disturbances in these pathways can have long-lasting reproductive consequences into adulthood. SV has been associated with increased androgen production in adult females, leading to hyperandrogenism and PMOS, and in some cases, reduced fertility (17, 33, 34). Consequently, there is concern that disturbances to follicular regulation during the prepubertal window could have lasting negative effects for long-term reproductive function and health. Our study demonstrated no detrimental impact of SV exposure on ovarian follicle number, health, or steroidogenesis within the prepubertal ovary, indicating that at least short-term exposure to SV may not adversely affect reproductive health in prepubertal females.

To date, no studies have examined the impact of prepubertal SV exposure on mouse ovaries *in vivo*. However, a previous rat study reported a marked reduction in follicle number and increased apoptotic markers following prepubertal SV treatment (35). Several methodological differences may explain the discrepancy between these findings and ours. Notably, Cansu *et al.* administered SV by oral gavage at 300 mg/kg/day for 24 days, a substantially longer exposure window than in this study. While oral administration reflects the clinical route, it is subject to first-pass hepatic metabolism and the reported serum SV concentrations were more than 1.5-fold below the lower limit of the therapeutic range. Our study administered SV intraperitoneally; this route does not allow direct administration to systemic circulation but does reduce the extent of first-pass hepatic metabolism. Taken together, the short-term exposure in this study did not infer ovarian damage, but a longer SV exposure may indeed have detrimental effects on the ovary. This is of clinical relevance, given that most patients prescribed SV as an antiepileptic would typically require long-term treatment with this drug.

Further evidence from adult rodent models highlights the importance of examining the impact of SV on different gonadal developmental stages. In adult female rats, 90-day administration of 400-600 mg/kg/day SV did not alter secondary follicle numbers, although ovarian cyst formation was observed (36). This aligns with clinical findings that adult females receiving SV treatment exhibit increased susceptibility to PMOS (17; 33). Given that our endpoint (PND17) precedes puberty and the onset of ovulation, and since ovarian cysts generally arise from follicles that fail to ovulate or release fluid (37), we did not expect to observe cysts at this stage. Future studies extending the SV exposure window from the prepubertal period into adulthood would be valuable to provide further insight into whether early-life SV treatment predisposes to PMOS-like phenotypes.

At the molecular level, we report no effect of SV on the expression of key steroidogenic genes involved in oestrogen and progesterone biosynthesis (CYP11A1, CYP19A1, STAR) in the prepubertal ovary, suggesting that short-term exposure does not impair ovarian steroidogenic capacity at this stage of development. This contrasts with an *in vitro* study in which SV was found to reduce CYP11A1 and STAR expression in granulosa-like tumour cell line (38). This discrepancy likely reflects differences in cell type, model systems, exposure duration, or species-specific sensitivity; to our knowledge, no prior *in vivo* study has examined steroidogenic gene expression in the prepubertal ovary.

This is the first investigation into the effect of SV on the prepubertal mouse testis. Prior studies have focused predominantly on fetal or adult stages, with variable outcomes. An *in vitro* study on the human fetal testis reported no effect of 72-hour SV exposure on germ cell density (19), whereas adult rodent models have demonstrated germ cell degeneration and reduced sperm counts (39–42). In our *in vivo* model, we observed a non-significant trend toward reduced germ cell density, which, if consistent with adult exposure data, suggests that SV-associated reproductive toxicity may be dose-dependent, with impact on germ cells influenced by testicular developmental stage. Given prior reports of SV-induced germ cell degeneration and reduced sperm counts in adult models, the observed trend towards decreased germ cell density reported here warrants careful consideration. If this effect is reproduced in the human testis, with a more prolonged exposure, it may carry implications for long-term spermatogenic capacity and male fertility in individuals treated with SV during childhood.

The absence of a fully established blood-testis-barrier (BTB) during the exposure window of the present study (PND6-10) is noteworthy. In the mouse, BTB formation occurs around PND10-16, meaning that during our exposure window the developing testis lacked the protective barrier that is present in the adult. The absence of the BTB prior to puberty has been proposed to render the prepubertal testis more susceptible to exogenous insults (43).

The lack of detrimental effect of SV on the developing testis shown here is therefore reassuring, since prepubertal SV exposure might have been anticipated to exert a more pronounced effect on the germ cells than adult exposure. Germ cells are essential for male fertility, giving rise to spermatozoa through spermatogenesis, a process sustained throughout life by a pool of SSCs. SSCs are established shortly after birth and constitute the self-renewing population that supports sperm production following pubertal HPG axis activation (44). During the prepubertal period, SSCs remain largely quiescent as the somatic testicular environment has not yet matured to support full spermatogenic differentiation (45). The preservation of SSC density across treatment groups is therefore also encouraging, indicating that the stem cell pool required to sustain spermatogenesis and support future fertility remains intact despite the overall trend toward reduced germ cell density.

Endocrine regulation is critical for SSC maintenance. Testosterone, secreted by Leydig cells, influences both SSC activity and Sertoli cell function (46). Low prepubertal testosterone levels help SSCs maintain a dormant state, whereas pubertal rise in testosterone initiates spermatogenesis (47). Importantly, SV has been shown to lower testosterone levels in male epilepsy patients (48) and in both *in vivo* and *ex vivo* models (19, 49). Endocrine-disrupting compounds may hinder androgen production through disruption of cholesterol transport, the expression of steroidogenic enzymes, or androgen receptor binding or action. Such alterations can result in incomplete masculinisation and malformations of the male reproductive tract, with subsequent impacts on fertility (50). Encouragingly, prepubertal SV exposure did not affect Sertoli and interstitial cell populations, nor expression of key steroidogenic genes in this study.

There are several limitations of the present study. Firstly, the exposure window was relatively short, comprising three doses administered over six days. SV is typically taken as a daily, long-term medication, and therefore it remains unclear whether more prolonged and/or cumulative exposure would affect gonadal health. Secondly, the doses employed may not have achieved therapeutically relevant serum concentrations, which limits direct comparison with clinical exposure scenarios. Finally, interspecies differences in drug metabolism and gonadal development may limit direct extrapolation to humans. Nonetheless, the mouse remains a well-established and ethically appropriate model for examining reproductive toxicology, with the present findings providing a foundation for more clinically translatable research, for example through the use of human *ex vivo* models or through cohort studies examining long-term reproductive outcomes in individuals treated with SV during childhood.

## CONCLUSION

The present study provides the first *in vivo* assessment of short-term prepubertal SV exposure on the developing reproductive system in both male and female mice. Ovarian follicle numbers and testicular architecture were found to be largely preserved, though a trend toward reduced germ cell density in males suggests that testicular germ cells may be more vulnerable than somatic cell populations. Taken together, these findings suggest that prepubertal exposure to SV may not impair gonadal development at the doses and timepoints examined. Future studies employing extended exposure protocols and longitudinal assessments extending into puberty and adulthood will be necessary to evaluate functional reproductive outcomes such as hormone production, sperm count, ovulation, and fertility. Such work will be essential to fully characterise the long-term reproductive risks associated with SV therapy during development and to inform evidence-based clinical guidelines for the management of epilepsy in paediatric patients.

## AUTHOR CONTRIBUTIONS

*Conception and design of the study:* RTM, KD and AS

*Acquisition and analysis of the data:* AL, AG, KD and AS

*Drafting of figures and manuscript:* AL

All authors critically reviewed and edited the article, as well as approving the final version of the manuscript.

## ACKNOWLEDGMENTS

We thank Prof. Norah Spears for support with the animal licence. We are grateful to Dr Grace Forsyth and Dr Sophie Thomson for their assistance with the RT-qPCR experiments. We also thank Adrian Garcia Burgos and Iain Porter at the IMPACT their support with imaging. We are grateful to Vivian Allison and Louise Dunn from the Histology laboratory for their assistance with histological processing. We also thank the animal facility technicians and the veterinarians, including Callum Davidson in particular, for their support with animal care and procedures and ethics.

## DATA AVAILABILITY STATEMENT

Data supporting the findings of this study are available from the corresponding author upon reasonable request.

