## Supplemental Table 1 for "Investigation into the effects of sodium valproate on prepubertal mouse gonads *in vivo*"

**Supplementary table 1**. List of antibodies used for immunofluorescence. Primary antibodies were selected to identify germ cells (MVH), Sertoli (SOX9) and interstitial cells (including Leydig cells; COUPTFII), and undifferentiated spermatogonia (PLZF). Species of origin, manufacturers, catalogue numbers, and working dilutions for both primary and fluorescent secondary antibodies are indicated.

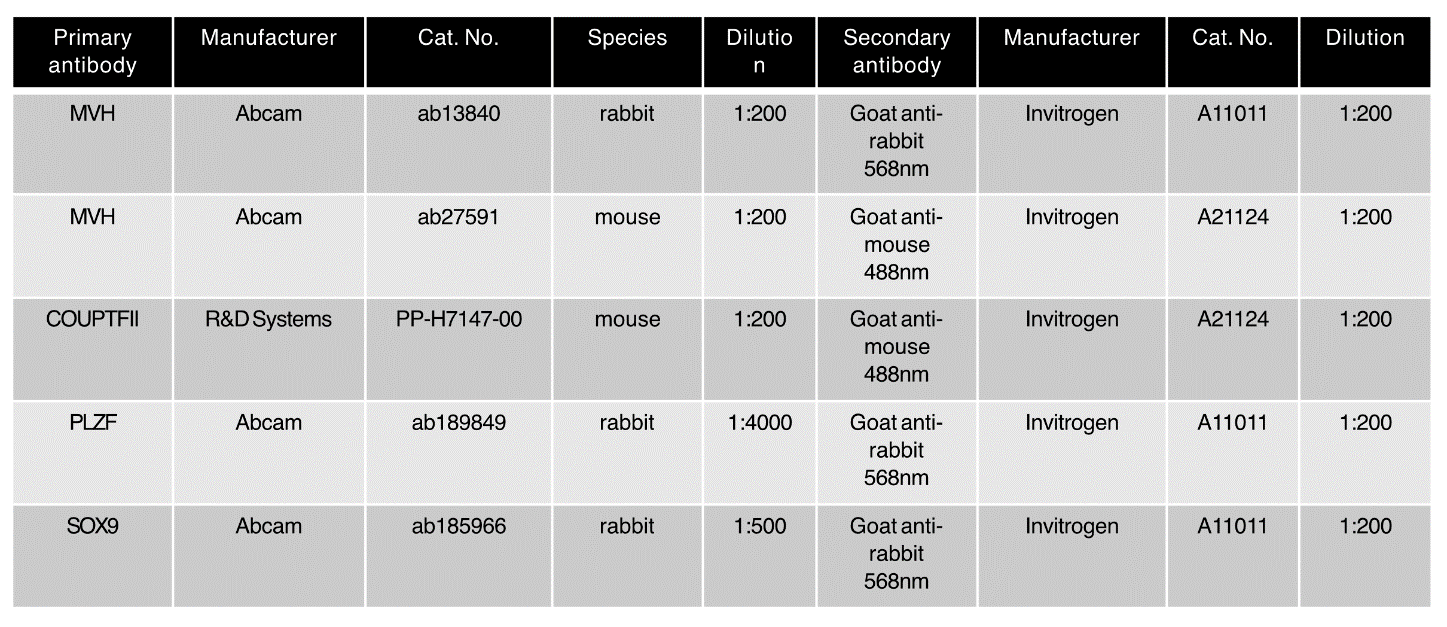
