## Supplemental Table 2 for "Investigation into the effects of sodium valproate on prepubertal mouse gonads *in vivo*"

**Supplementary table 2.** List of primers used. Genes were selected to assess housekeeping gene expression (GAPDH, ACTB), steroidogenic capacity (STAR, CYP11A1), Leydig cell function in testes (INSL3), and oestrogen biosynthesis in ovaries (CYP19A1). Final concentration of primers used was 200nM. All primers were manufactured by Integrated DNA Technologies Inc.

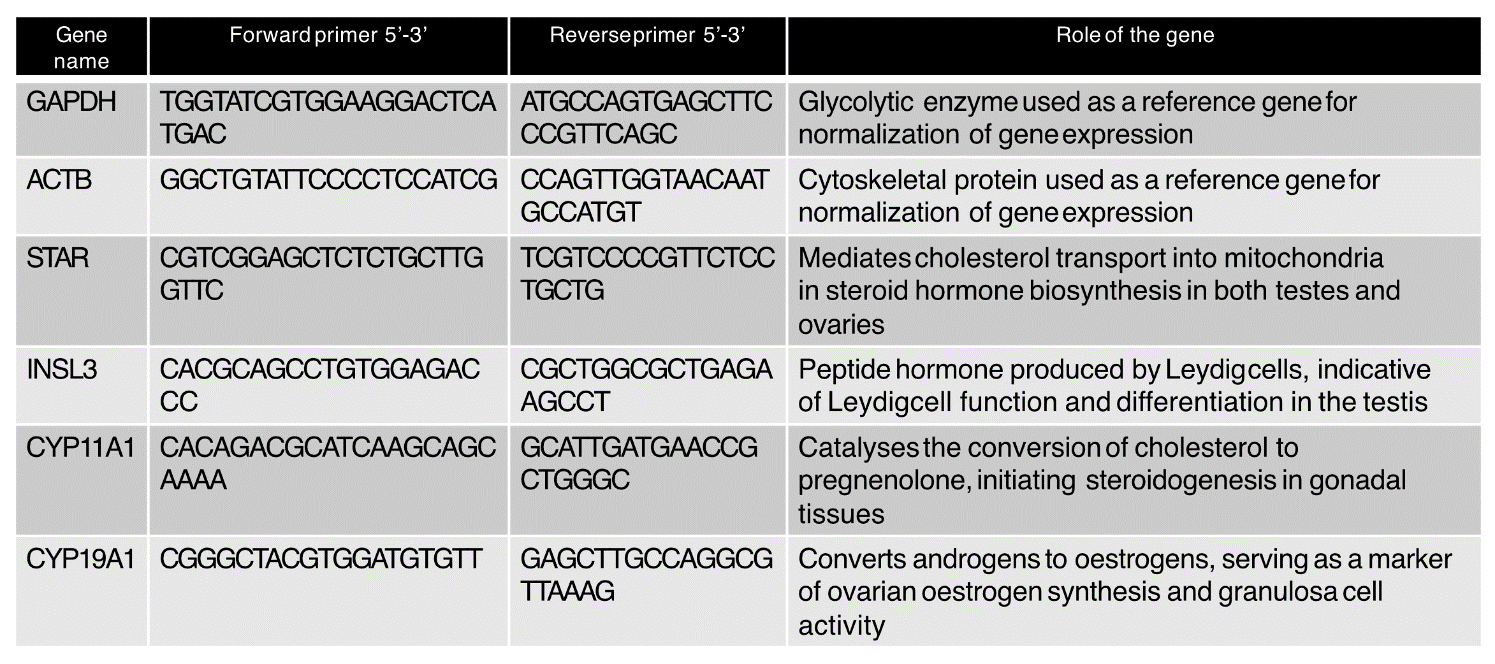
