## Supplemental Figure 1 for "Investigation into the effects of sodium valproate on prepubertal mouse gonads *in vivo*"

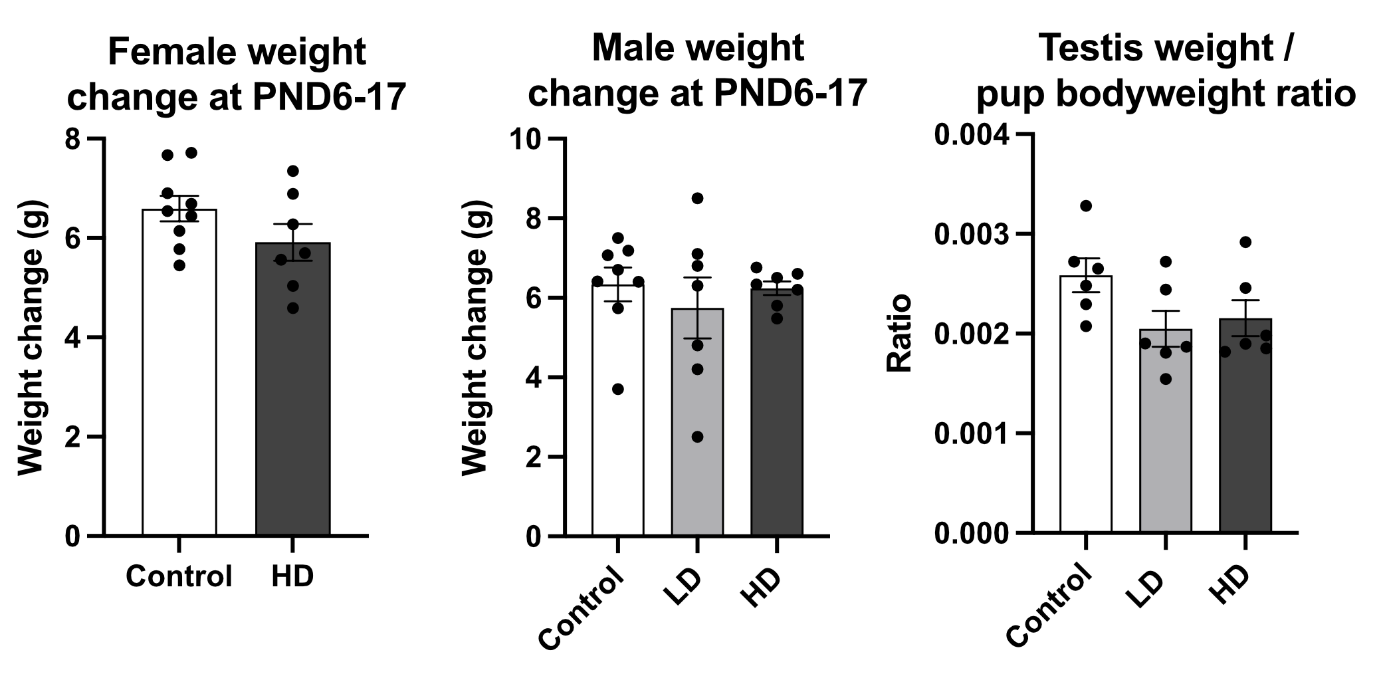

**Supplementary Figure 1.** Bodyweight and testis-to-bodyweight ratio of pups. Female (A) and male (B) pup bodyweight and testis weight-to-bodyweight ratio (C) were unchanged after low dose (LD, 50 mg/kg) or high dose (HD, 100 mg/kg) SV treatment (PND6–PND17). Bars represent mean ± SEM.
