## Supplemental Figure 2 for "Investigation into the effects of sodium valproate on prepubertal mouse gonads *in vivo*"

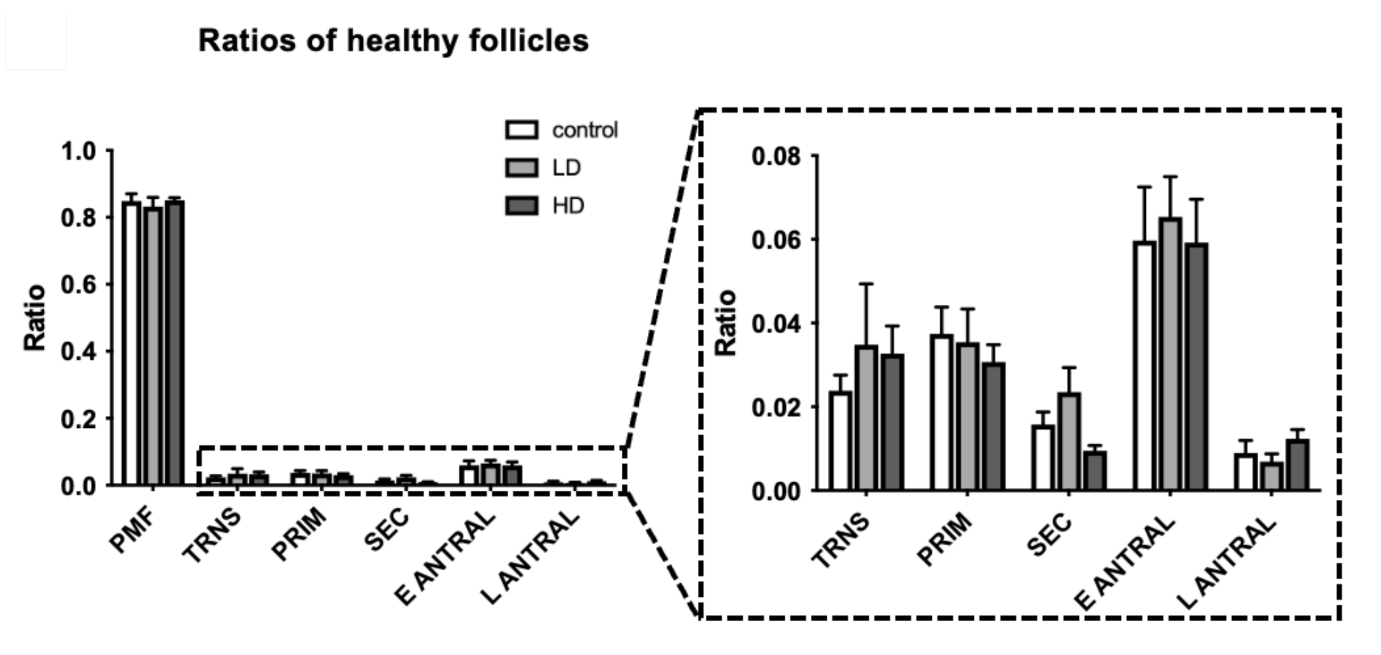

**Supplementary Figure 2.** Ratios of healthy follicle types in prepubertal ovaries. Ratios of primordial (PMF), transitional (TRNS), primary (PRIM), secondary (SEC), early antral (E ANTRAL), and late antral (L ANTRAL) follicles relative to the total number of healthy follicles in prepubertal mouse ovaries at PND17 following saline (control), low dose (LD; 50 mg/kg), or high dose (HD; 100 mg/kg) SV exposure. Ratios were calculated as the number of follicles of a given developmental stage divided by the total number of healthy follicles per ovary. Bars represent mean + SEM; n = 6 per group.
